# Tubulin isotypes contribute opposing properties to balance anaphase spindle morphogenesis

**DOI:** 10.1101/2025.04.04.647290

**Authors:** Emmanuel T. Nsamba, Abesh Bera, Vaishali Todi, Landon Savoy, Ryan M. Gupta, Mohan L. Gupta

## Abstract

Faithful chromosome segregation requires proper function of the mitotic spindle, which is built from, and depends on, the coordinated regulation of many microtubules and the activities of molecular motors and MAPs. In addition, microtubules themselves are assembled from multiple variants, or isotypes of α- and β-tubulin, yet whether they mediate the activities of motors and MAPs required for proper spindle function remains poorly understood. Here, we use budding yeast to reveal that α-tubulin isotypes regulate opposing outward- and inward-directed forces in the spindle midzone that facilitate optimal spindle elongation and length control. Moreover, we show that the isotypes mediate balanced spindle forces by differentially localizing the antagonistic force generators Cin8 (kinesin-5) and Kar3 (kinesin-14) to interpolar microtubules. Our results reveal new roles for tubulin isotypes in orchestrating motor and MAP activities and provide insights into how forces in the spindle are properly calibrated to ensure proper mitotic spindle morphogenesis.

## Introduction

The mitotic spindle is vital for cell replication and genomic stability. It is a dynamic structure built largely from microtubules, which are intrinsically dynamic cytoskeletal filaments polymerized from tubulin, a heterodimer of α- and β-tubulin subunits. Mitotic spindles are composed of three classes of microtubules, differing in arrangement, dynamic behavior and function, yet assembled from a common pool of tubulin (Dumont and Mitchison, 2009; Guilloux and Gibeaux, 2020). One class, astral microtubules, extend outward from the spindle poles and facilitate spindle positioning. The other two classes reach inwards toward the central spindle: interpolar microtubules establish crosslinks with those from the opposite pole to form an antiparallel, overlapping bundle called the midzone, and kinetochore microtubules connect the spindle poles to the sister chromatids. Spindle assembly, morphogenesis, and overall function can only be achieved via the coordinated action of these distinct microtubule populations, orchestrated by a range of motor and non-motor regulators acting as nucleators, polymerizers, depolymerizers, crosslinkers, stabilizers, destabilizers, as well as force generators. Post-translational modification (PTM) of tubulin can also influence some aspects of spindle properties (reviewed in (Lopes and Maiato, 2020; Janke and Magiera, 2020; Akera, 2023)). Yet, how distinct microtubule subpopulations are discretely and timely regulated to control spindle morphogenesis remains essentially unknown.

The spindle continuously changes throughout mitosis while maintaining the structural integrity required for chromosome segregation (Rizk et al., 2014; Thomas et al., 2020). Microtubule motor proteins as well as non-motor microtubule-associated proteins (MAPs) located at various sites in the spindle and at the cell cortex mediate spindle dynamics (Goshima and Scholey, 2010; Guilloux and Gibeaux, 2020). In the midzone, where interpolar microtubules from opposite poles are crosslinked by MAPs, motor proteins generate sliding forces on the antiparallel microtubules that contribute to spindle elongation and size control (Dumont and Mitchison, 2009). For example, conserved kinesin-5 motors localize to the midzone and support bipolar structure by pushing spindle poles apart, while kinesin-14 motors exert inward-directed forces that balance the activity of kinesin-5. In the budding yeast, *S. cerevisiae*, the kinesin-5, Cin8, plays a major role in sorting interpolar microtubules into antiparallel bundles during spindle assembly and uses plus-end directed motility to slide them apart during anaphase (Saunders et al., 1997b; Leary et al., 2019). These activities are antagonized by the kinesin-14, Kar3 (Saunders et al., 1997b), which also crosslinks spindle microtubules and uses minus-end directed motility to align antiparallel microtubules and generate inward forces to regulate spindle length (Hepperla et al., 2014; Saunders et al., 1997b; a). The proper balance between Cin8 and Kar3 is essential. Spindle morphogenesis is impaired without Cin8 (Hoyt et al., 1992) or Kar3 (Saunders et al., 1997a; Meluh and Rose, 1990), but also by excess of either activity (Saunders et al., 1997b). Similarly, chromosome instability is increased in the absence of either Cin8 (Yuen et al., 2007) or Kar3 (Yuen et al., 2007; Zhu et al., 2015), yet also by excess Cin8 (Ouspenski et al., 1999) or Kar3 (Duffy et al., 2016). How the activities of these major regulators are balanced to produce healthy spindles is unknown.

In addition to microtubule regulators and PTMs, in most eukaryotes microtubules are made from multiple variants, or isotypes, of α- and β-tubulin subunits. Relative to regulatory factors and PTMs, the role of tubulin isotypes in controlling microtubule functions is poorly understood (Reviewed in (Nsamba and Gupta, 2022)). The presence of specific isotypes can be important for aspects of spermatogenesis (Hutchens et al., 1997; Hoyle and Raff, 1990), oocyte maturation (Feng et al., 2016), cilia function (Hurd et al., 2010; Silva et al., 2017), blood platelet formation (Strassel et al., 2019; Schwer et al., 2001), and the nervous system (Lockhead et al., 2016; Zheng et al., 2017; Baran et al., 2010; Bittermann et al., 2019; Buscaglia et al., 2020; Latremoliere et al., 2018). However, the underlying mechanisms remain obscure. Moreover, whether tubulin isotypes regulate microtubules themselves, motors or MAPs within the mitotic spindle is completely unknown.

Using budding yeast we reveal that tubulin isotypes regulate opposing outward- and inward-directed forces in the spindle midzone that facilitate optimal spindle elongation and length control. Moreover, we show that the isotypes mediate balanced spindle forces by differentially localizing the antagonistic force generators Cin8 (kinesin-5) and Kar3 (kinesin-14) to interpolar microtubules. Overall, our results provide the first mechanistic evidence for the role of tubulin isotypes in proper mitotic spindle morphogenesis.

## Results

### Tubulin isotypes differentially affect mitotic spindle function

The mitotic spindle in most eukaryotes, including yeast, is composed of three major classes of microtubules: kinetochore, interpolar, and astral (Fig. 1a). Although they are assembled from the same pool of tubulin subunits, each class must be differentially regulated to ensure the spindle is fully functional (Reviewed in (Guilloux and Gibeaux, 2020)). To test whether tubulin isotypes differentially mediate interpolar microtubule function we monitored spindles in cells harboring solely Tub1 or Tub3 at levels comparable to total α-tubulin in normal cells (named Tub1-only and Tub3-only, respectively). To visualize microtubules we used an otherwise identical, exogenous GFP-Tub1 or GFP-Tub3 construct in Tub1- or Tub3-only cells, respectively. Corresponding control cells harboring both Tub1 and Tub3 were similarly visualized with an exogenous copy of GFP-Tub1 or GFP-Tub3, denoted WT-Tub1 or WT-Tub3. To achieve efficient chromosome segregation, anaphase spindle length must, to some extent, scale with cell size (Rizk et al., 2014; Goshima and Scholey, 2010; Guilloux and Gibeaux, 2020). We found, however, that anaphase spindle length and/or morphology is not properly regulated in Tub1-only cells. Relative to control, Tub1-only cells display an over 5-fold increase in fishhook spindles. In sharp contrast, Tub3-only cells are reduced 87% compared to control cells (Fig. 1b-c).

**Figure 1.**
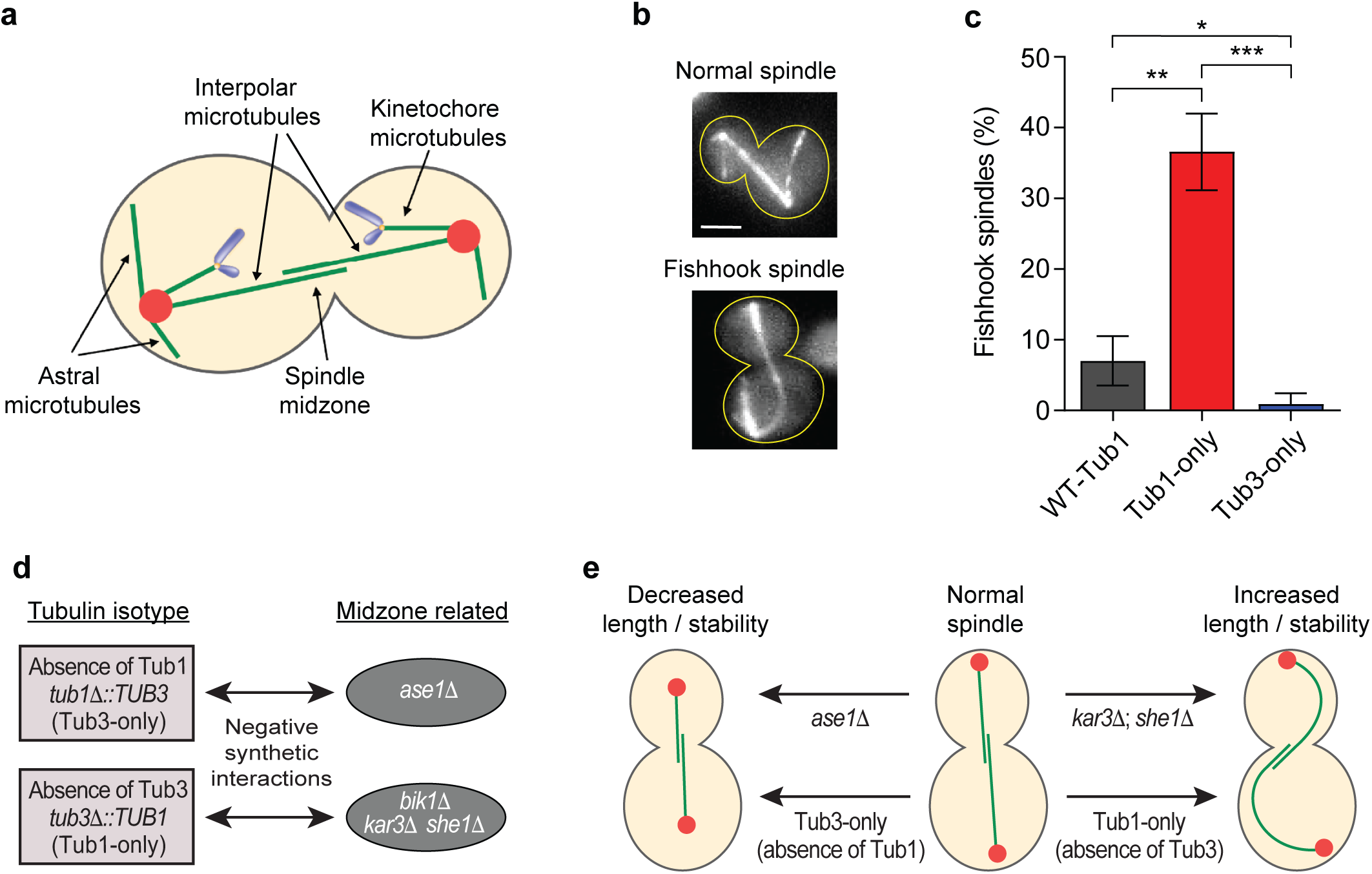
Tubulin isotypes differentially affect mitotic spindle function. (**a**) Schematic of yeast mitotic spindle with three microtubule populations (green), chromosomes (purple) and Spindle Pole Bodies (SPBs; red). (**b**) Representative images of normal (straight) and fishhook (bent) spindle morphology in cells expressing GFP-tubulin. Scale bar = 2 μm. (**c**) Percentage of anaphase spindles with fishhook morphology during late anaphase. For WT-Tub1 vs. Tub1-only or Tub3-only, p = 0.014 and 0.05, respectively, and for Tub1-only vs. Tub3-only p = 0.0004. Mean ± SD of three trials with 50 spindles per genotype for trials 1 and 2, and 44, 38, and 37 spindles for WT-Tub1, Tub1-only and Tub3-only, respectively in trial 3. (**d**) Negative genetic interactions between the absence of *TUB1* (top; Tub3-only; *tub1Δ::TUB3*) or *TUB3* (bottom; Tub1-only; *tub3Δ::TUB1*) and loss of midzone-related proteins identified by Synthetic Genetic Array (SGA) screening (Nsamba et al., 2021). (**e**) Relationship of spindle phenotypes in Tub1-only, Tub3-only and cells lacking spindle midzone proteins identified by SGA in (**d**).

Fishhook spindles, which elongate beyond the length of the cell and bend within cellular constraints, can result from impaired spindle disassembly (Woodruff et al., 2010), upregulated dynein activity (Woodruff et al., 2009), or defects in spindle length control (Rizk et al., 2014). Thus, the absence of Tub3 likely results in one or more of these defects. Previous work indeed reported increased dynein activity on astral microtubules in Tub1-only cells (Nsamba et al., 2021). To gain insight into whether the tubulin isotypes also differentially mediate interpolar microtubule function we examined the negative genetic interactions between midzone-related components and either α-tubulin isotype (Nsamba et al., 2021). Negative genetic interactions are characterized by greater fitness impairment due to the simultaneous perturbation of two genes than the predicted additive effect of each perturbation alone. In the extreme, such interactions can result in synthetic lethality but useful information is also gained from interactions that are less than lethal, or synthetic sick compared to the single perturbations (Tong et al., 2001). Moreover, large scale analysis shows genes with similar function are likely to display negative genetic interactions (Tong et al., 2004).

Notably, the absence of Tub1 or Tub3 produces distinct negative genetic interactions with midzone-related proteins (Fig. 1d). This relationship suggests the α-tubulin isotypes contribute different properties to the mitotic spindle via the function of interpolar microtubules. The absence of Tub3 (Tub1-only) displays negative genetic interactions with She1, Kar3, or Bik1. She1 stabilizes metaphase spindles as a microtubule crosslinker (Zhu et al., 2017), but also destabilizes spindles following anaphase (Woodruff et al., 2010). Kar3 also contributes a destabilizing activity and/or inward forces to the anaphase spindle (Saunders et al., 1997b; Cottingham et al., 1999). Tub3 may, therefore, potentially contribute similar properties to the spindle. On the other hand, absence of Tub1 (Tub3-only) is synthetic sick with the loss of Ase1, that normally functions to stabilize the midzone and facilitate spindle elongation (Janson et al., 2007; Schuyler et al., 2003). Thus, Tub1 may also function to promote spindle stability and/or elongation.

### Loss of Tub3 enhances bipolar spindle assembly and elongation

To elucidate the roles of Tub1 and Tub3 in spindle morphology, we examined bipolar spindle dynamics, a process that depends on antiparallel crosslinking and sliding of interpolar microtubules (Guilloux and Gibeaux, 2020). Cells with spindle pole bodies (SPBs) tagged with RUBY (Cnm67-mRuby2) and GFP-tagged microtubules were released from G1 arrest (alpha-factor) and imaged at 15-min intervals. We found that bipolar spindles form significantly sooner in cells utilizing solely Tub1 (Fig. 2a-b). This was evident in both the first and second mitotic cycles following synchronized release (i.e. 45 and 105 min). In addition to bipolar structure, spindles in Tub1-only cells deliver a pole (SPB) into the bud compartment sooner than control cells (Fig. 2c-d), which became significant by the second synchronized cycle (i.e. 60 and 120 min).

**Figure 2.**
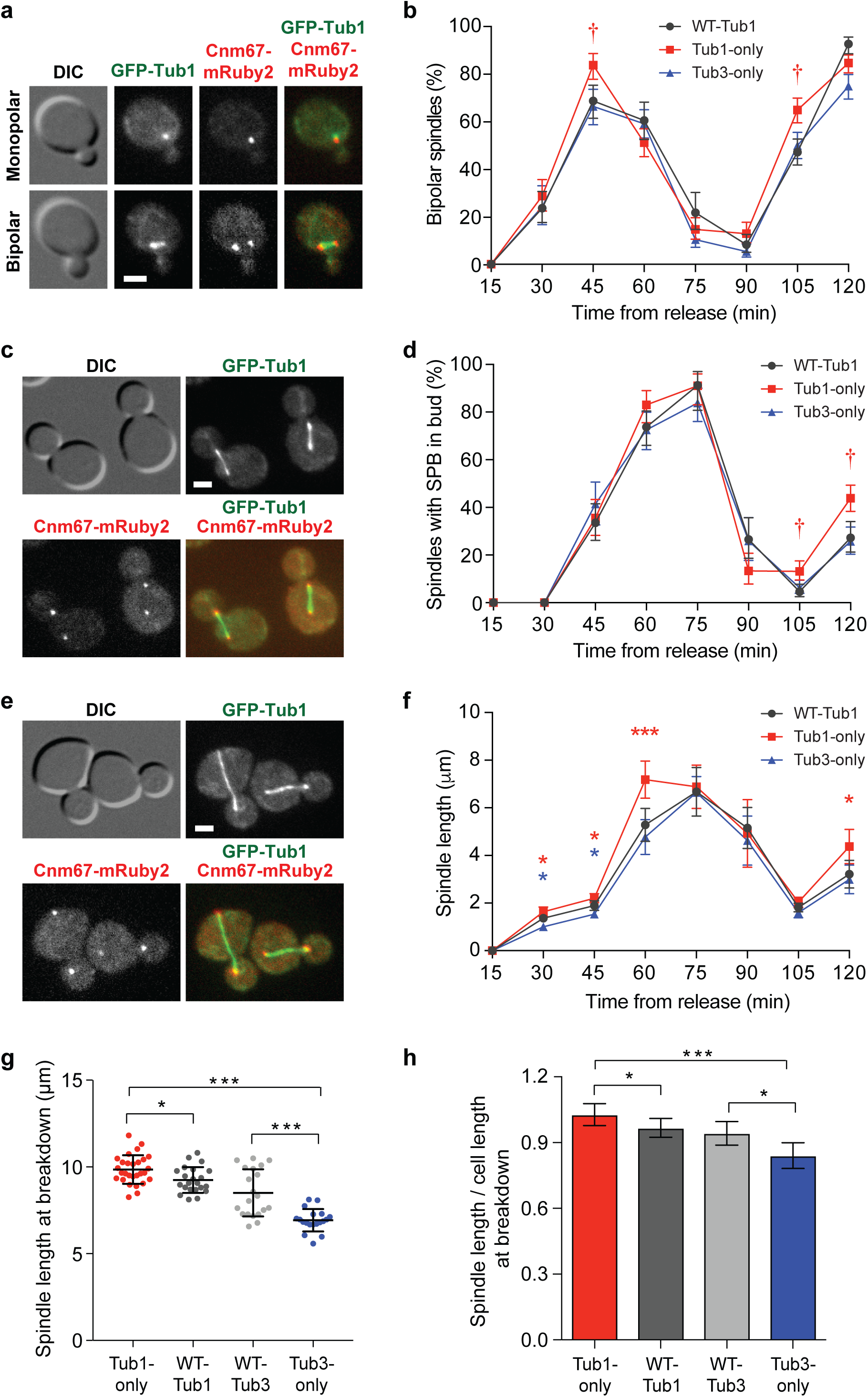
Loss of Tub3 enhances bipolar spindle assembly and elongation. (**a**) Representative DIC and wide-field fluorescence images of GFP-Tub1 (microtubules) and Cnm67-mRUBY2 (SPBs) of a WT-Tub1 cell with a monopolar or bipolar spindle. (**b**) Percent cells with bipolar spindle following release from G1 arrest in cells of the indicated genotypes. (**c**) Representative images of WT-Tub1 cells with a bipolar spindle in the mother (right) or in the bud (left) compartment. (**d**) Percent cells with an SPB in the bud following release from G1 arrest. (**e**) Representative images of WT-Tub1 cells with varying spindle lengths. (**f**) Average spindle length following release from G1 arrest. (**g**) Anaphase spindle length and (**h**) length relative to cell length at breakdown in cells of the indicated genotypes. Compared to spindles in control cells, those in Tub1-only and Tub3-only cells are significantly longer and shorter, respectively. In (**b**) and (**d**) error bars represent 95% confidence interval from error of proportion; †significant difference (p < 0.05) for Tub1-only vs. WT-Tub1 based on confidence intervals. In (**f**) error bars represent SEM; *p ≤ 0.05, ***p ≤ 0.001 for the indicated color vs. WT-Tub1. For (**b**) n for 15 min = 33, 49, 51, for 30 min = 116, 178, 192, for 45 min = 166, 181, 195, for 60 min = 278, 159, 296, for 75 min = 289, 120, 253, for 90 min = 300, 230, 256, for 105 min = 337, 340, 340, and for 120 min = 293, 250, 385 for Tub3-only, WT-Tub1, and Tub1-only cells, respectively, from three trials. For (**d**) n for 30 min = 116, 178, 192, for 45 min = 123, 155, 166, for 60 min = 135, 156, 130, for 75 min = 118, 57, 89, for 90 min = 100, 113, 120, for 105 min = 334, 299, 296, and for 120 min = 241, 198, 331 for Tub3-only, WT-Tub1, and Tub1-only cells, respectively, from three trials. For (**f**) n for 30 min = 80, 80, 80, for 45 min = 60, 51, 60, for 60 min = 42, 40, 42, for 75 min = 45, 31, 38, for 90 min = 22, 21, 22, for 105 min = 59, 53, 60, and for 120 min = 58, 60, 60 for Tub3-only, WT-Tub1, and Tub1-only cells, respectively, from two trials. For (**g, h**) bars or lines show mean ± SD. For Tub1-only, WT-Tub1, WT-Tub3, Tub3-only in (**g**) n = 28, 22, 20, 20 and in (**h**) n = 44, 31, 34, 32, respectively, from three independent days. For all panels *p ≤ 0.05, ***p ≤ 0.001. Only statistically significant comparisons are indicated. Scale bars = 2μm.

Spindles in Tub1-only cells could enter the bud sooner due to increased dynein-mediated pulling on astral microtubules, which is more efficient on Tub1-microtubules (Nsamba et al., 2021). Accelerated SPB delivery may also stem from Tub1-only cells entering anaphase sooner and/or Tub1-only spindles elongating faster than those in control cells. A combination of these mechanisms could also be involved. Consistent with the latter two possibilities, the average spindle length increases earlier in Tub1-only compared to control cells (Fig. 2e-f). While in Tub3-only cells the average spindle length lags that in control cells (Fig. 2e-f).

We next monitored spindle morphogenesis using time-lapse microscopy. Spindles in Tub3-only cells fail to reach normal lengths and are disassembled at a maximum length of 6.4 ± 0.6 µm compared to 7.9 ± 1.4 µm in control (WT-Tub3) cells (Fig. 2g; p < 0.001). Consistent with disassembly before attaining normal length, Tub3-only spindles reach only 84 % of the cell length compared to 94 % in control cells (Fig. 2h; p < 0.05). In contrast, Tub1-only spindles are longer relative to controls (WT-Tub1), at 9.3 ± 0.8 vs. 8.7 ± 0.7 µm, respectively (Fig. 2g; p < 0.05), and often exceed cell length at disassembly (Fig. 2h; 102 % vs. 97 %, respectively; p < 0.05). Notably, spindles in Tub1-only cells are 46% longer than those in Tub3-only cells at the time of breakdown (Fig. 2g; p < 0.001). Altogether, these data suggest Tub1 facilitates spindle elongation and Tub3 functions to antagonize it.

### Tub1 and Tub3 differentially localize motors that control anaphase spindle elongation

Shorter or fishhook spindles, as well as slower or faster elongation may result from imbalances in the inward and outward spindle forces or perturbed midzone stability. The combined genetic and phenotypic data suggest Tub1 promotes, and Tub3 antagonizes spindle stability and/or elongation. Yeast spindles exhibit a biphasic anaphase B, with an initial fast phase followed by a second slower phase (Yeh et al., 1995; Kahana et al., 1995). To test the molecular basis for tubulin isotype function exclusively on interpolar spindle microtubules, we examined the localization of midzone-associated MAPs and molecular motors.

Loss of the antiparallel microtubule crosslinker and midzone stabilizing factor, Ase1, is synthetic sick with the absence of Tub1 (Tub3-only; Fig. 1d) (Nsamba et al., 2021). Although Ase1 localization to Tub1-only, relative to Tub3-only spindles, is higher in both fast and slow phases of anaphase, this increase does not reach statistical significance (Supplementary Fig. S1a-d). The plus-end tracking protein Bim1, which localizes to the spindle and contributes to midzone stability (Woodruff et al., 2010; Gardner et al., 2008), also does not show statistically significant differences between Tub1- and Tub3-only spindles during the fast or slow phases (Supplementary Fig. S1e-g). Thus the observed spindle defects are likely not due to altered MAP localization within the spindle midzone.

In contrast, localization of the kinesin-5, Cin8, which crosslinks midzone microtubules and generates outward forces to elongate the spindle is significantly enhanced on Tub1-only compared to Tub3-only spindles (Fig. 3a-b). This is particularly evident during the initial, fast phase of anaphase in which Cin8 is enriched 1.4-fold on Tub1-only spindles (Fig. 3c; p ≤ 0.01). During the slow phase Cin8 is increased 22% (Supplementary Fig. S1h; p = 0.087). This relationship is maintained when spindles are binned by length, where pre-anaphase spindles (1-2 μm) and those in slow phase (6-8 μm) are indistinguishable and significant differences are seen throughout the fast phase (3-6 μm; Fig. 3d).

**Figure 3.**
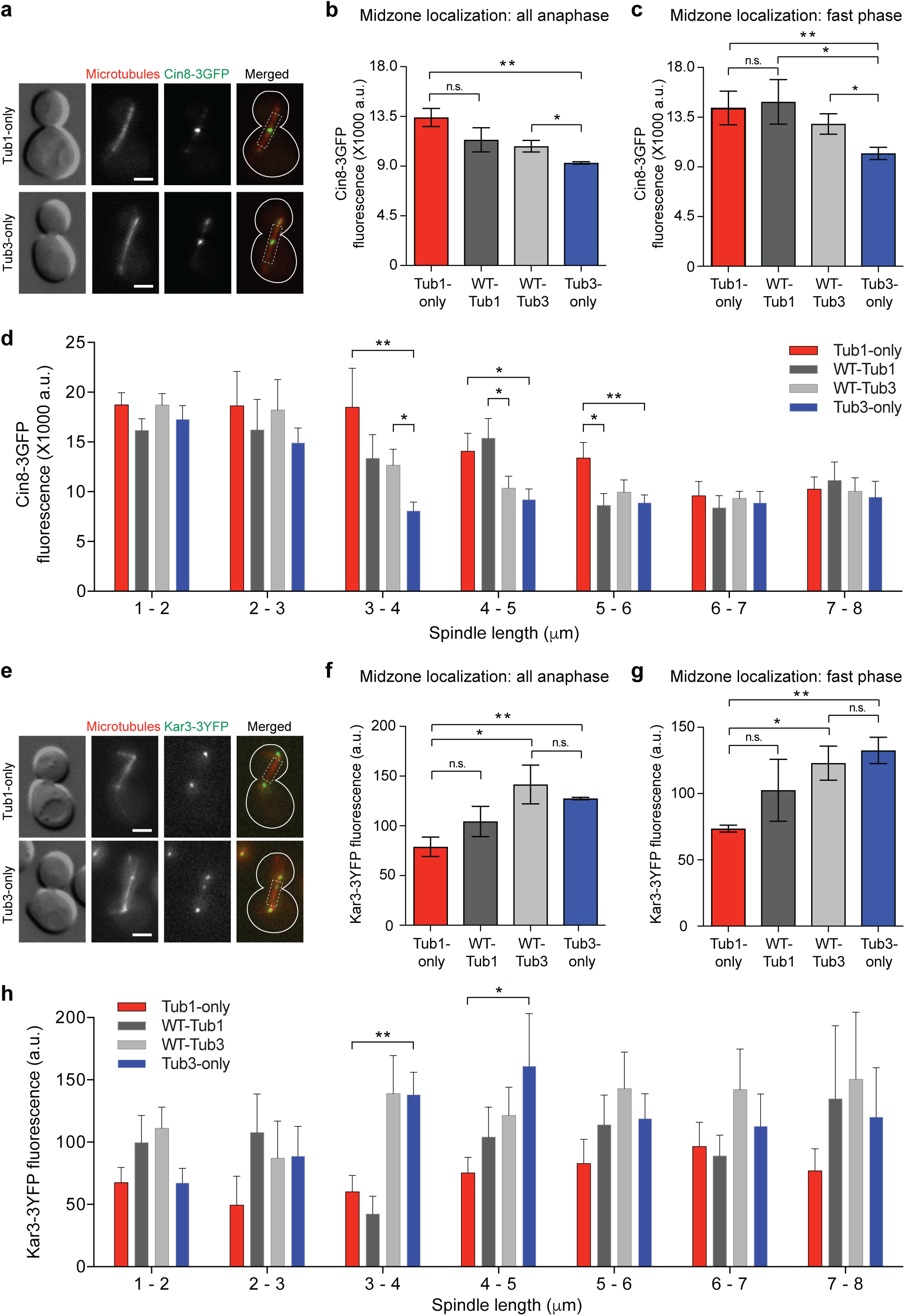
Tub1 and Tub3 differentially localize motors that control anaphase spindle elongation. (**a-d**) Cin8 localization to the spindle. (**a**) Representative DIC and wide-field fluorescence images of Cin8-3GFP localization to the spindle (microtubules) in Tub1-only (top) and Tub3-only (bottom) cells. Hashed box in merged image represents area of spindle quantified. (**b-d**) Cin8-3GFP fluorescence signal on spindles (**b**) in all anaphase cells, (**c**) during the initial, fast phase of anaphase, and (**d**) on spindles binned according to length in cells of the indicated genotypes. (**e-h**) Kar3 localization to the spindle. (**e**) Representative DIC and wide-field fluorescence images of Kar3-3YFP localization to the spindle (microtubules) in Tub1-only (top) and Tub3-only (bottom) cells. Hashed box in merged image represents the region over which the line-scan between SPBs was quantified. (**f-h**) Kar3-3YFP fluorescence signal on spindles (**f**) in all anaphase cells, (**g**) during the initial, fast phase of anaphase, and (**h**) on spindles binned according to length in cells of the indicated genotypes. For (**a, e**) microtubules are visualized by mRuby3-Tub1 or mRuby3-Tub3; merged images show microtubules in red and YFP/GFP-tagged proteins in green. In graphs bars represent mean ± SEM from three trials; in (**b**) n for Tub1-only = 10, 20, 21, WT-Tub1 = 13, 10, 20, WT-Tub3 = 5, 8, 5, Tub3-only = 10, 13, 17; in (**c**) n for Tub1-only = 30, 41, 40, WT-Tub1 = 37, 33, 40, WT-Tub3 = 35, 42, 31, Tub3-only = 41, 39, 50; in (**d**) for 1-2μm n = 60, 61, 51, 32, 2-3μm n = 12, 15, 9, 20, 3-4μm n = 12, 20, 17, 21, 4-5μm n = 35, 25, 30, 46, 5-6μm n = 32, 29, 44, 27, 6-7μm n = 28x, 39, 44, 27, 7-8μm n = 24, 15, 23, 8 for Tub1-only, WT-Tub1, WT-Tub3, and Tub3-only, respectively; in (**f**) n for Tub1-only = 14, 11, 10, WT-Tub1 = 13, 7, 10, WT-Tub3 = 11, 7, 12, Tub3-only = 12, 7, 8; in (**g**) n for Tub1-only = 25, 22, 20, WT-Tub1 = 23, 23, 20, WT-Tub3 = 21, 17, 31, Tub3-only = 24, 23, 18; in (**h**) for 1-2μm n = 17, 11, 12, 6, 2-3μm n = 6, 5, 10, 9, 3-4μm n = 10, 5, 12, 17, 4-5μm n = 15, 17, 15, 9, 5-6μm n = 11, 11, 10, 17, 6-7μm n = 11, 12, 11, 9, 7-8μm n = 10, 8, 6, 6 for Tub1-only, WT-Tub1, WT-Tub3, and Tub3-only, respectively. For all panels *p ≤ 0.05, **p ≤ 0.01, n.s. = not significant. All comparisons not marked are n.s. Scale bars = 2 μm.

Loss of Kar3, which antagonizes the activity of Cin8 in the spindle (Saunders et al., 1997b), displays a strong negative genetic interaction with the absence of Tub3 (Tub1-only; Fig. 1d) (Nsamba et al., 2021). Kar3 contributes destabilizing activity and/or inward forces to the spindle, and its loss also displays a strong negative interaction with loss of Cin8 (Costanzo et al., 2016). Together these data indicate that Kar3 opposes the function of Cin8 in the spindle midzone and that Tub3 may promote spindle activities similar to those of Kar3. Strikingly, Kar3 localization is increased >50% on Tub3-only relative to Tub1-only spindles (Fig. 3e-f). This difference is most pronounced during fast anaphase, when Kar3 localization is nearly 2-fold higher on Tub3-only relative to Tub1-only spindles (Fig. 3g, p ≤ 0.01; Supplementary Fig. S1i). Like Cin8, pre-anaphase spindles (1-2 μm) and those in slow phase (6-8 μm) show insignificant differences, with significantly more Kar3 on Tub3-only spindles during the fast phase (Fig. 3h). Overall these results suggest Cin8 function is enhanced by Tub1 and in contrast Kar3 function is enhanced by Tub3.

### Tub1 and Tub3 are opposing regulators of anaphase spindle elongation

The increased Cin8 and decreased Kar3 on Tub1-only spindles predicts they elongate at a faster rate. Conversely, the decreased Cin8 and increased Kar3 on Tub3-only spindles suggests they display slower elongation. We thus monitored spindle elongation by timelapse imaging (Fig 4a). In control cells, the fast elongation phase proceeded at 0.58 and 0.46 µm/min in WT-Tub1 and WT-Tub3 cells, respectively (Fig. 4b-c). Compared to these intermediate rates of control spindles, the fast phase was accelerated in Tub1-only and reduced in Tub3-only cells (Fig. 4b-c). Strikingly, the elongation rate was reduced nearly 50% in Tub3-only relative to Tub1-only cells (0.35 relative to 0.64 µm/min; Fig. 4c). During the subsequent slow phase, spindle elongation is less affected but remains significantly reduced in Tub3-only relative to Tub1-only cells (0.13 vs. 0.17 µm/min; Fig. 4c). In general, spindle elongation rates in WT-Tub1 and WT-Tub3 controls, which harbor both tubulin isotypes, are intermediate relative to those in single isotype cells (Fig. 4c).

**Figure 4.**
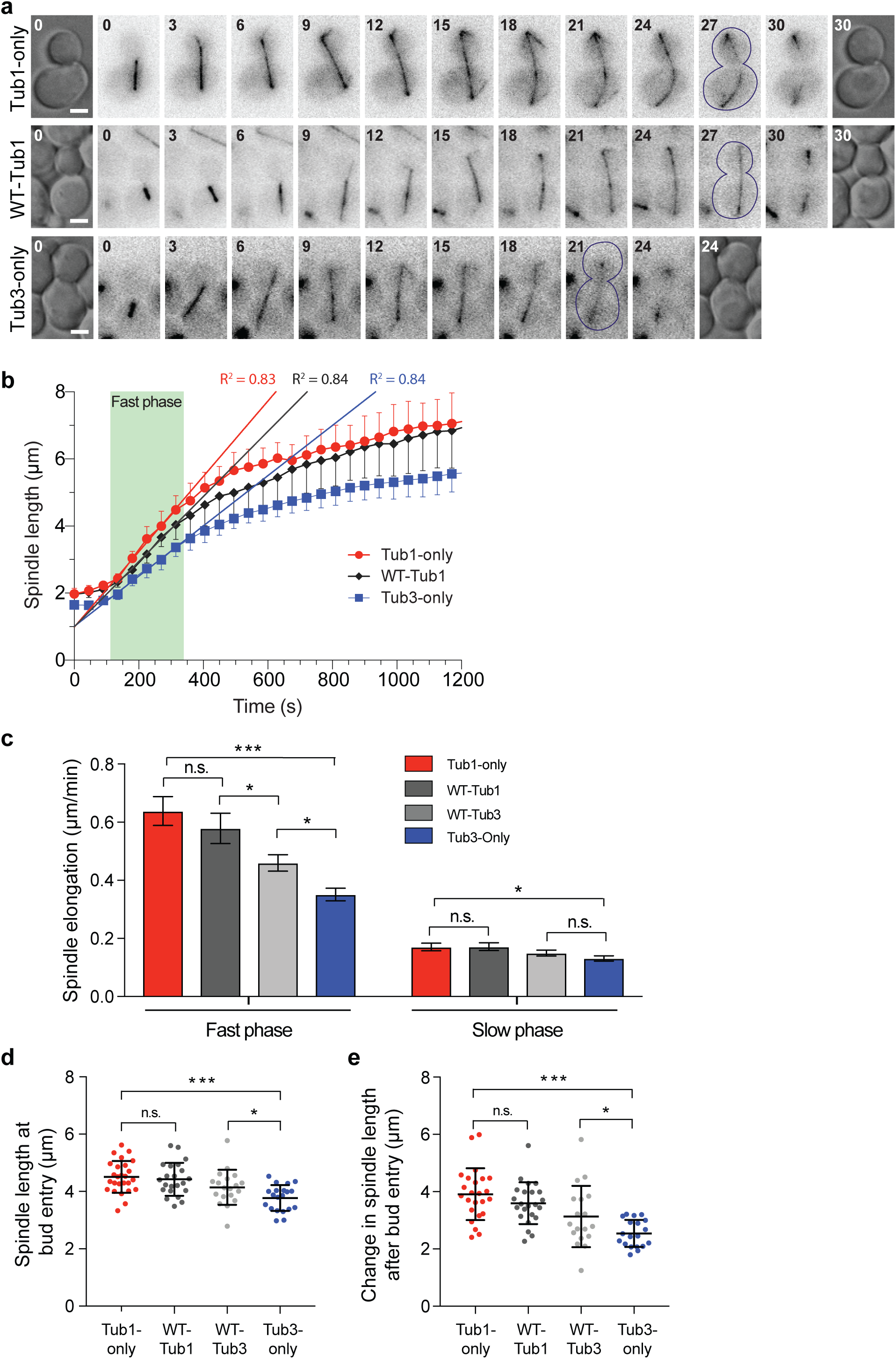
Tub1 and Tub3 are opposing regulators of anaphase spindle elongation. (**a**) Representative timelapse images of spindle elongation showing fishhook morphology in Tub1-only (top) and limited elongation in Tub3-only (bottom) cells. DIC image of cell is at beginning and end. Fluorescence images of microtubules are shown with inverse intensity (GFP-Tub1 for Tub1-only and WT-Tub1; GFP-Tub3 for Tub3-only). Time is shown in minutes. (**b**) Average spindle length over time during anaphase in cells of the indicated genotypes. Lines show linear regression through the fast phase. (**c**) Average anaphase spindle elongation rates during the initial fast and subsequent slower phase. (**d**) Spindle length at time of SPB bud entry. (**e**) Spindle elongation from time of SPB bud entry to spindle breakdown. (**b-e**) Graphs show mean ± SD. For (**b**) n = 10 for each genotype; for Tub1-only, WT-Tub1, WT-Tub3, Tub3-only in (**c**) n = 18, 18, 17, 22 for fast and 18, 19, 17, 22 for slow phase; in (**d**) n = 26, 22, 19, 20 and in (**e**) n = 24, 23, 18, 18, respectively, from three independent trials. For all panels *p ≤ 0.05, ***p ≤ 0.001, n.s. = not significant. Scale bars = 2 μm.

Previous work demonstrated that the two major spindle positioning mechanisms are differentially supported by Tub1 and Tub3 (Nsamba et al., 2021). Thus, it is possible that the spindle elongation rate may be indirectly constrained by the mother cell cortex or SPB passage through the bud neck due to impairment of the early or late positioning mechanisms, respectively. To test this possibility, we monitored changes in anaphase spindle length within the mother cell and after bud entry. Tub1-only spindles elongated significantly more than Tub3-only spindles in the mother cell prior to bud entry (4.5 vs. 3.7 μm; p < 0.001; Fig. 4d). Moreover, Tub1-only spindles continued elongating significantly more than Tub3-spindles after an SPB had entered the bud compartment (3.9 vs. 2.5 μm; p < 0.001; Fig. 4e). The extent of spindle elongation in either compartment of control cells, expressing both isotypes, is intermediate compared to Tub1- and Tub3-only cells (Fig. 4d-e).

Thus, independent of the roles of Tub1 and Tub3 in differentially supporting Dyn1- and Kar9-mediated spindle positioning mechanisms, respectively (Nsamba et al., 2021), Tub1-only spindles elongate faster during anaphase, and to a greater extent than control spindles harboring both Tub1 and Tub3. Conversely, Tub3-only spindles elongate slower and to a lesser extent than control spindles.

### Differences in spindle elongation do not result from changes in dynein activity

Dynein motor activity contributes outward pulling forces on SPBs via astral microtubules during anaphase (Yeh et al., 1995; Adames and Cooper, 2000). Thus, it is possible that the increased dynein function in Tub1-only cells (Nsamba et al., 2021) may contribute to their increased spindle elongation rate (Fig. 4c). Similarly, the decreased dynein function in Tub3-only cells (Nsamba et al., 2021) could explain their slower spindle elongation and lack of fishhook spindles. To test this possibility, we monitored spindle elongation rates in cells lacking dynein (*dyn1Δ*). If dynein activity is responsible for the increased spindle elongation rate in Tub1-only cells then its loss would suppress this phenotype. In contrast, we found that elongation rates are generally similar whether or not cells possess dynein (compare Fig. 4c and Fig. 5a). Notably, relative to control *dyn1Δ* cells harboring both tubulin isotypes, spindles in Tub1-only *dyn1Δ* cells continue to elongate significantly faster, while those in Tub3-only *dyn1Δ* cells elongate significantly slower (Fig. 5a). As with cells harboring functional dynein (Fig. 4c), these differences were most pronounced during the fast phase of anaphase (Fig. 5a).

**Figure 5.**
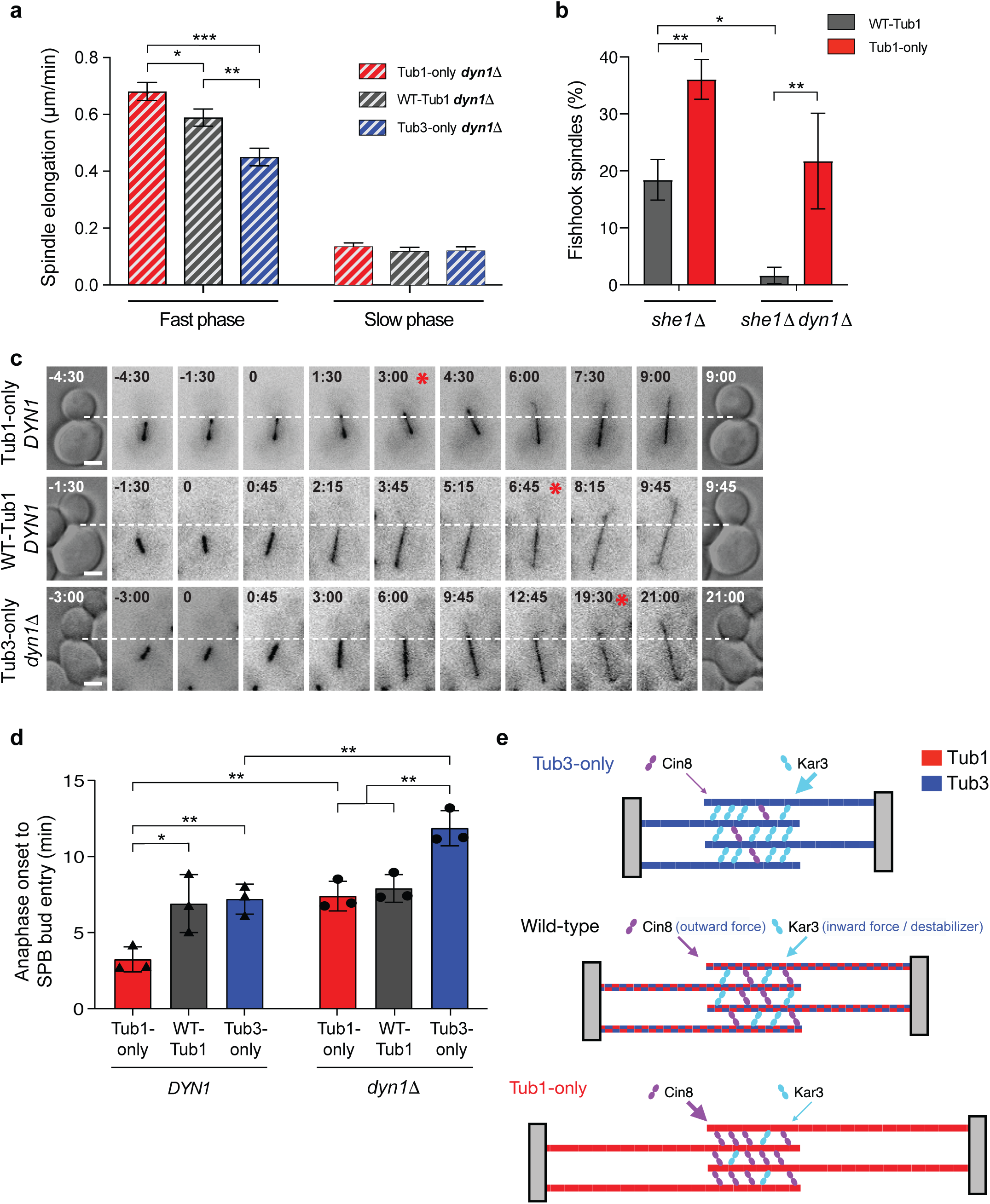
Differences in spindle elongation are independent of dynein activity, but dynein mediates timing of anaphase spindle positioning. (**a**) Average anaphase spindle elongation rates in cells lacking dynein (*dyn1Δ*) during the initial fast and subsequent slower phase. (**b**) Percentage of anaphase spindles with fishhook morphology in control and Tub1-only cells lacking She1 (left; *she1Δ*) or lacking both She1 and dynein (right; *she1Δ dyn1Δ*). (**c**) Representative timelapse images of anaphase onset to SPB bud entry for the indicated genotypes. DIC image of cell is at beginning and end and fluorescence images of microtubules are shown with inverse intensity (GFP-Tub1 for Tub1-only and WT-Tub1; GFP-Tub3 for Tub3-only). Hashed line represents position of bud neck, time relative to anaphase onset is shown in minutes, red asterisks indicate SPB bud entry, scale bars = 2 μm. (**d**) Average time from anaphase onset to SPB bud entry in cells harboring (left) or lacking (right) dynein activity. (**e**) Model of tubulin isotype function in spindle elongation. Spindles assembled exclusively from the α-tubulin isotype Tub3 (top; blue) accumulate more Kar3 and less Cin8, resulting in delayed and slower elongation, and thus subsequently shorter spindles that are not correctly scaled to cell length at the time of disassembly. Spindles composed exclusively of Tub1 (bottom; red) preferentially bind more Cin8 than Kar3, leading to higher elongation rates and excessive elongation prior to disassembly. In wildtype cells (middle), Cin8 and Kar3 are balanced by the presence of both Tub1 and Tub3, thus spindle elongation functions normally and spindle length is appropriately scaled to cell length. In (**a**) for Tub1-only, WT-Tub1, Tub3-only (all *dyn1Δ*), n = 38, 29, 18 for fast and 31, 30, 18 for slow phase, respectively; in (**b**) for WT-Tub1 *she1Δ*, Tub1-only *she1Δ*, WT-Tub1 *she1Δ dyn1Δ*, Tub1-only *she1Δ dyn1Δ* n = 50, 43, 43, 41 in trial one, 18, 15, 11, 8 in trial two, and 33, 40, 46, 47 in trial three, respectively; in (**d**) bar represents average of three trials (symbols) where n = 19, 17, 23, 11, 9, 11 in trial one, 18, 23, 25, 10, 9, 8 in trial two, and 8, 18, 15, 5, 6, 3 in trial three for Tub3-only, WT-Tub1, Tub1-only, Tub3-only *dyn1Δ*, WT-Tub1 *dyn1Δ*, Tub1-only *dyn1Δ*, respectively. Graphs shows mean ± SD; *p ≤ 0.05, **p ≤ 0.01, ***p ≤ 0.001.

The fishhook phenotype generally results from spindle length exceeding cell length, hence forcing the spindle to adopt a bent conformation (Straight et al., 1998; Rizk et al., 2014). It is also possible that bent spindles could be facilitated by dynein-mediated pulling on astral microtubules lateral to the spindle axis. Thus, we considered whether the increased fishhooks in Tub1-only cells could result from increased dynein activity. To test this possibility, we first quantified fishhook spindles in cells with increased dynein by removing its inhibitor, She1 (Markus et al., 2012). In control cells, loss of She1 significantly increases fishhook frequency from 7% to ∼19% (Fig. 1c and Fig. 5b). Notably, the fishhook frequency in Tub1-only *she1Δ* cells remains double that in *she1Δ* control cells (Fig. 5b). The fishhook percentage in Tub1-only *she1Δ* cells also remains comparable to the level in Tub1-only cells harboring functional She1, at ∼36% (compare Fig. 1c and Fig. 5b). The fact that *she1Δ* does not increase fishhooks in Tub1-only cells is consistent with dynein being already highly active and/or efficient in these cells.

Moreover, the finding that fishhooks in Tub1-only cells, whether in the presence or absence of She1, are nearly double than in control *she1Δ* cells, which have hyperactive dynein, is indicative of an alternative mechanism, such as increased outward midzone forces, contributing to the excessive fishhooks in Tub1-only cells. To verify this, we quantified fishhooks in *dyn1Δ* cells, which lack any lateral-directed pulling on astral microtubules. In control *she1Δ dyn1Δ* cells, fishhooks drop to less than 2%, indicating nearly all fishhooks require dynein activity to manifest in normal cells (Fig. 5b). In Tub1-only *she1Δ* cells that also lack Dyn1, the fishhook frequency remains ∼22% (Fig. 5b). This confirms that approximately half the fishhooks in *she1Δ* Tub1-only cells occur independent of dynein-mediated pulling on astral microtubules, and are most likely due to excessive midzone pushing forces.

Altogether these data show that the increased spindle elongation rate and fishhook outcome in Tub1-only cells do not result from dynein pulling forces on astral microtubules, but instead from additional robust outward pushing forces generated within the spindle midzone. Furthermore, the slower spindle elongation in Tub3-only does not result from decreased dynein pulling on astral microtubules, but rather from lower pushing forces and/or increased opposing inward forces generated in the midzone.

### Dynein-mediated pulling forces are needed for Tub3-only, but accelerate Tub1-only anaphase spindle positioning

We previously showed that Tub3 enhances Kar9-mediated spindle positioning which moves the spindle near and aligns it with the bud neck prior to anaphase onset, and that during anaphase, Tub1 optimizes the dynein-mediated mechanism that pulls an SPB through the neck (Nsamba et al., 2021). Independent of spindle positioning, we now find that Tub3 inhibits, while Tub1 enhances spindle elongation via midzone pushing forces. Thus, we considered whether the changes in anaphase spindle elongation rates could potentially contribute to their effects on spindle positioning. To test this idea, we monitored the time required from anaphase onset to deliver an SPB into the bud or, in other words, successful spindle positioning (Fig. 5c).

In control cells harboring both Tub1 and Tub3 it takes on average 6.9 min from anaphase unset until an SPB is delivered to the bud (Fig. 5d). The result is similar in Tub3-only cells (7.2 min) where pre-anaphase spindles are closer and better aligned with the neck, indicating that some combination of optimal pre-anaphase positioning with decreased dynein and spindle elongation is sufficient to achieve normal timing of anaphase positioning (Fig. 5d). By sharp contrast, in Tub1-only cells, which are impaired for pre-anaphase spindle positioning, an SPB is delivered to the bud in less than half the time (3.2 min; p < 0.05; Fig. 5d). This outcome could be due to enhanced dynein pulling forces, increased outward spindle pushing forces, or a combination of the two.

To discriminate these possibilities, we monitored the time for SPB bud delivery in cells lacking dynein, excluding those spindles that were sufficiently mispositioned that they failed to enter the bud. In control cells lacking dynein, SPB delivery into the bud is slightly but not significantly delayed compared to cells with function dynein (7.9 vs. 6.9 min; p = 0.46; Fig. 5d). This demonstrates that a combination of normal pre-anaphase positioning and anaphase elongation can facilitate SPB delivery into the bud independent of dynein. Strikingly, when Tub1-only cells lack dynein, SPB entry is significantly delayed (7.4 vs 3.2 min; p = <0.01; Fig. 5d). Notably, this timing is indistinguishable from control *dyn1Δ* cells that harbor both Tub1 and Tub3 (7.4 min vs. 7.9 min; p = 0.55; Fig. 5d). Thus, the accelerated SPB entry in Tub1-only cells is largely a result of enhanced dynein function. Without dynein, the impaired pre-anaphase positioning in the absence of Tub3 (Nsamba et al., 2021) may be compensated by then enhanced spindle elongation forces in Tub1-only cells (Fig. 5d). When Tub3-only cells also lack dynein, SPB delivery into the bud during anaphase is significantly delayed relative to control cells lacking dynein (11.9 vs. 7.9 min; p < 0.01; Fig. 5d). Despite enhanced Kar9-mediated pre-anaphase spindle positioning (Nsamba et al., 2021), Tub3-only cells are reliant on dynein function for timely SPB delivery to the bud during anaphase (11.9 vs. 7.2 min for *dyn1Δ* vs. *DYN1*, respectively; p < 0.01; Fig. 5d). Thus, while spindles in Tub3-only cells lack sufficient elongation forces to efficiently push an SPB through the bud neck in the absence of dynein, the combination of enhanced dynein function and spindle elongation accelerates SPB delivery in Tub1-only cells.

## Discussion

Our findings provide novel insights into the roles of tubulin isotypes in mediating spindle morphogenesis. Most eukaryotic organisms harbor multiple isotypes of α- and β-tubulin that can copolymerize into microtubules, yet in most cases their contributions to general or specialized microtubule functions has remained largely elusive. Here, we used budding yeast to interrogate the role of tubulin isotypes in spindle morphogenesis. During spindle assembly, interpolar microtubules emanating from opposite spindle poles are crosslinked in an antiparallel manner to form overlapping bundles (Janson et al., 2007). Microtubules in these bundles then grow and are slid past each other, pushing the spindle poles apart and contributing to physical chromosome separation in anaphase (Masuda et al., 1988; Pavin and Tolić, 2015). Our data reveal that this process is regulated by the α-tubulin isotypes (Fig. 5e). *TUB1* and *TUB3* exhibit distinct, negative genetic interactions with spindle stabilizers and destabilizers, respectively, which suggests the isotypes contribute unique and opposing properties to effective spindle function.

Consistent with this premise, functional assays revealed mechanistic insights into how Tub1 and Tub3 regulate spindle morphology during anaphase. During the fast phase of anaphase, Tub1 preferentially localizes the midzone stabilizer and outward force generating kinesin-5 motor, Cin8, to interpolar microtubules, while diminishing localization of the antagonistic kinesin-14 motor, Kar3 (Fig. 3). This is consistently significant between spindles composed exclusively of Tub1 or Tub3 throughout the fast phase, demonstrating the impact of either isotype. Significant differences in Cin8 localization can also be seen with control strains at certain spindle lengths. It should be noted, however, that fluorescence labeling of microtubules requires an exogenous copy of tagged tubulin. Thus, the WT-Tub1 and WT-Tub3 strains contain proportionally more of their corresponding isotype than an unlabeled wildtype cell. Although such an unlabeled control could potentially be developed using non-tubulin labeling, the results show Cin8 and Kar3 localization are generally proportional to the Tub1/Tub3 gene ratio. Moreover, spindle localization data with Tub1- and Tub3-only cells contrast the total absence of either isotype. Excess Cin8 promotes premature and excessive spindle elongation, while surplus Kar3 produces the opposite effect (Saunders et al., 1997b). Analogously, increased Tub1 causes more rapid, and excessive spindle elongation (Fig. 1c, Fig. 4, Fig. 5b). As opposed to Tub1-only spindles, those made exclusively of Tub3 preferentially localize Kar3 and inhibit recruitment of the antagonistic motor Cin8 during fast anaphase (Fig. 3). In accordance, spindles built with extra Tub3 elongate at a significantly slower rate and fail to reach normal length during anaphase (Fig. 4, Fig. 5e). The differences in elongation rates are largest during the initial phase of anaphase, when the role of Cin8 is most prominent (Straight et al., 1998). Budding yeast utilize two kinesin-5 family members. While Cin8 is required for the fast phase, cells lacking Kip1 have defects in the slow phase of anaphase spindle elongation (Straight et al., 1998). Although the most prominent isotype-dependent effects on spindle elongation occur during the fast phase, the difference in the slow phase between Tub1-only and Tub3-only is statistically significant (Fig. 4c). Thus, it will be informative to determine whether Kip1 is also differentially localized to Tub1- over Tub3-only spindles, particularly during late anaphase.

Whereas Tub3-only spindles fail to reach the length observed in control cells, spindles in Tub1-only cells do not properly limit spindle elongation. As a result, fishhook spindles are nearly absent in Tub3-only cells, but over 5-fold more prevalent in Tub1-only compared to control cells (Fig. 1c). The percentage of fishhooks increases nearly 3-fold in control cells lacking the dynein inhibitor, She1 (Woodruff et al., 2009), demonstrating that dynein pulling on astral microtubules can facilitate fishhook formation (Fig. 5b). The fact that fishhooks do not increase in Tub1-only cells upon She1 loss (Fig. 1c vs. Fig. 5b) is consistent with dynein function being optimized by Tub1 (Nsamba et al., 2021). When dynein activity is removed from these cells (*dyn1Δ*), the fishhook frequency in those harboring the combination of both isotypes drops to below 2% (Fig. 5b), indicating that dynein-mediated pulling on astral microtubules is the main mechanism of fishhook initiation in normal cells. However, in Tub1-only cells lacking dynein the fishhook percentage remains 13-fold higher than comparable control cells, and over half that in Tub1-only cells with active dynein. Revealing an alternate mechanism, Tub1-only spindles fail to terminate anaphase elongation and exceed cell length due, as our data indicate, to excessive midzone pushing forces. Conversely, the lack of fishhook spindles in Tub3-only cells is consistent with increased Kar3, decreased Cin8, and ultimately, diminished outward forces generated in the midzones of Tub3-only spindles.

Altogether, these data reveal that tubulin isotypes impart distinct properties to microtubules that specifically regulate spindle elongation and overall length during anaphase. The preferential localization of Cin8 and Kar3, by Tub1 and Tub3, respectively, serves to calibrate the activities of these opposing regulators to properly balance the forces controlling spindle morphogenesis (Fig. 5e). Consistent with this premise, spindles in control cells harboring both isotypes exhibit intermediate phenotypes to those harboring exclusively Tub1 or Tub3. In addition to influencing localization, Tub1 and/or Tub3 may potentially possess characteristics that specifically accommodate the molecular mechanisms used by Cin8 and Kar3, respectively. Besides molecular motors, our data show the spindle stabilizing proteins, Bim1 (Woodruff et al., 2010; Gardner et al., 2008) and Ase1 (Thomas et al., 2020), are similarly localized to both Tub1- and Tub3-only spindles, which reinforces our model that antagonistic motors are the major driver of the isotype-mediated morphological changes during anaphase. Although their localization is essentially comparable, however, we cannot exclude the possibility that the activities of Bim1, Ase1, or other spindle-associated factors are differentially influenced by interactions with either isotype. Despite this possibility, it remains clear that Tub1 and Tub3 impart distinct, and opposing, properties to interpolar microtubules and the morphological function of the mitotic spindle.

The function of Tub1 and Tub3 in mediating opposing spindle activities is reminiscent of their roles in differentially optimizing the major spindle positioning mechanisms (Nsamba et al., 2021). Analogous to their functions in the spindle midzone revealed here, Tub3 preferentially localizes components of the Kar9 pathway to astral microtubules and optimizes the function of this pre-anaphase spindle positioning mechanism. Conversely, Tub1 promotes localization of proteins in the dynein pathway to astral microtubules and enhances this anaphase spindle positioning mechanism. Together these studies provide compelling evidence that tubulin isotypes provide unique qualities that allow microtubules assembled from multiple isotypes to efficiently perform the range of required cellular functions. Previous studies have shown that α- and β-tubulin isotypes are important for mediating specialized microtubule-mediated functions in microorganisms (reviewed in (Bera and Gupta, 2022)) and in higher eukaryotes (reviewed in (Nsamba and Gupta, 2022)). For example, in *D. melanogaster* specific isotypes are needed during spermatogenesis (Hutchens et al., 1997; Hoyle and Raff, 1990). While in *C. elegans* certain isotypes influence neurite outgrowth (Lockhead et al., 2016; Zheng et al., 2017; Baran et al., 2010) and proper cilia function (Hurd et al., 2010; Silva et al., 2017). In mouse (*M. musculus*) individual isotypes are known to function in blood platelet formation (Strassel et al., 2019; Schwer et al., 2001) as well as neuronal development (Bittermann et al., 2019; Buscaglia et al., 2020) and axon regeneration (Latremoliere et al., 2018). Additionally, the activity of MAPs (Boscheron et al., 2016; Denarier et al., 2021), kinesins (Silva et al., 2017; Sirajuddin et al., 2014), and depolymerases (Ti et al., 2018) can be influenced by the isotype composition of microtubules. In essentially all cases, however, the underlying mechanisms remain obscure. Altogether these data support our finding, that copolymerized isotypes differentially support, or optimize, multiple microtubule-dependent mechanisms, is likely a highly conserved characteristic across tubulin isotypes. Elucidating the underlying molecular mechanism(s) will be critical to enable direct analysis of how Tub1 and Tub3, or other isotypes alter the binding and/or biophysical properties of molecular motors and/or MAPs. With spindle morphogenesis, for example, localization of motors to midzone microtubules may be mediated directly or indirectly involving other proteins or complexes, and may be dependent on midzone-specific microtubule structures and/or factors/complexes. We are currently pursuing such experiments in the lab.

Collectively these studies suggest a model in which individual tubulin isotypes, although constrained by the requirements to support dynamic instability and serve as an adequate platform for general microtubule functions, can independently adapt to better facilitate the mechanisms deployed by divergent MAPs, motors, and regulatory factors. Using a single α- and β-isotype would restrict all microtubule-interacting factors in the cell to conform to an identical binding interface and molecular dynamics properties. If any factor sufficiently diverged it would fail to adequately interact with or regulate microtubule function. Likewise, if the lone α- or β-tubulin isotype were to adapt to a divergent or novel regulator, it may perturb the activities of existing factors. In budding yeast, the two α-tubulin isotypes differ in their abilities to support distinct microtubule functions, and the loss of either disrupts the balance needed to properly perform microtubule-dependent processes. The presence of many α- and β-tubulin variants can facilitate the mechanisms of a wider range of interacting and regulatory factors required to support the diversity of microtubule functions in complex organisms. Likewise, mutations in individual isotypes may preferentially disrupt the activities of specific MAPs, motors, and/or regulators, and compromise distinct processes required for proper development and health.

## Materials and Methods

### Yeast strains and plasmids

Yeast strains used in this study are derivatives of the S288C background and, together with plasmids, are listed in Supplemental Table S1. Development of Tub1-only (*tub3Δ*::*TUB1*) and Tub3-only (*tub1Δ*::*TUB3*) strains together with GFP- and mRUBY3-tubulin plasmid constructs was described in (Nsamba et al., 2021). Microtubules were marked using an exogenous copy of α-tubulin as reported (Straight et al., 1997). Spindle pole bodies were visualized using Cnm67-Ruby2 (Gazy et al., 2013) introduced by genetic cross. Previously reported constructs were used to tag Cin8-3GFP (Aiken et al., 2014), Ase1-3GFP (Zhu et al., 2017), and Bim1-GFP (Liakopoulos et al., 2003; Nsamba et al., 2021) at their endogenous loci. For Kar3, the integrating LEU2 plasmid, pMG100, harboring the coding sequence for the C-terminal region of Kar3 fused in-frame with 3YFP was linearized by cutting within the Kar3 sequence with NheI and used to tag *KAR3* at the endogenous locus. Deletion of genes such as *dyn1Δ*::*TRP1* was achieved by fragment-mediated homologous recombination. For functional analyses and assays, tagged and/or deleted alleles were first introduced and verified in control strains and then crossed into Tub1- and Tub3-only strains to generate mutant and equivalent control strains. For genetic crossing and tetrad analysis, complete tetrads were confirmed by verifying proper segregation of the genetic markers as well as mating types.

### Analysis of spindle assembly and morphology during cell cycle progression

To induce G1 arrest yeast cultures were grown to early log phase (A_600_ = ∼0.4) in rich media at 30°C, then treated with 10 μM α-factor (Zymo Research, Y1001) for 2.5 – 3 h. Microscopic analysis confirmed > 95% of cells were arrested. The α-factor was then washed out and cells released into fresh media at 24°C with samples fixed every 15 min by incubation in 3.7% formaldehyde for 15 minutes at room temperature followed by washing three times in PBS buffer. Cells were imaged as described below.

Spindles were scored as bipolar by the presence of two Cnm67-Ruby2-tagged spindle poles (SPBs) with GFP-tubulin signal between them. Spindle length and the presence of one spindle pole in the daughter cell were determined as previously reported (Aiken et al., 2014) using SlideBook software (Intelligent Imaging Innovations; RRID:SCR_014423). Fishhook spindles were defined as those that exceeded cell length and displayed a bent morphology.

### Live cell microscopy

Cells were imaged as previously reported (Nsamba et al., 2021). Briefly, cultures grown overnight in synthetic complete (SC) medium supplemented with 300 μg/ml adenine were diluted and grown an additional ∼6 h (over two doublings) at 24°C to reach OD_600_ ∼ 0.3. Cells were mounted on 1.2% agarose pads also containing SC + adenine as described previously (Luchniak et al., 2013). Imaging was performed at 24°C unless otherwise stated. Exposure times were optimized depending on the duration of the experiment, the protein of interest and the fluorescent label. Automated 3D image acquisition was conducted using an AxioImager M2 microscope (Carl Zeiss) with a 63X 1.4 NA Plan-APOCHROMAT oil immersion objective, Semrock filter sets (Semrock), piezoelectric-driven Z-stage, and CoolSNAP HQ^2^ charged-coupled device (CCD) camera (Photometrics) driven by SlideBook software. Image analysis was performed using SlideBook and ImageJ (NIH; RRID:SCR_003070).

### Spindle elongation analysis

Z-series timelapse images of live cells expressing GFP-tubulin were collected every 45 s for 90 min. Spindle length was determined in three dimensions (i.e., across z-planes when appropriate). Because SPBs occasionally move toward each other concomitant with midzone disengagement the length at the time of spindle breakdown was scored using the time point 90 s before breakdown. The periods of initial fast and subsequent slow phases of anaphase spindle elongation were identified as previously reported (Thomas et al., 2020). Linear regression analysis identified the following start and end lengths for the initial fast phase of anaphase; Tub1-only (2.43 – 5.45 μm), WT-Tub1 (2.14 – 5.06 μm), WT-Tub3 (2.11 – 4.34 μm), and Tub3-only (2.01 – 4.3 μm). The slow phase of anaphase encompassed length changes following the fast phase until spindle breakdown. The rates of spindle elongation in each phase were determined by linear regression through the linear portion of each phase, lasting at least 180 s.

The time from anaphase onset until SPB bud entry was calculated from the time a pre-anaphase spindle (i.e. 1 – 2 μm) began sustained elongation until one SPB passed through the neck into the bud compartment.

### Fluorescence intensity analysis of tagged proteins

The spindle localization of GFP/YFP tagged proteins on mRUBY3 labeled microtubules was measured essentially as described previously (Thomas et al., 2020). Two-color images were recorded using 10 z-planes spaced 0.75 μm apart, and only cells with the entire spindle in focus, determined by the visibility of both SPBs, were used for analysis. The z-series images were sum projected, and spindle associated signal was measured in a region along the spindle encompassing the protein of interest, excluding the region and any signal peaks associated with the two SPBs, with a width of at least four pixels. Spindle associated signal within the region was determined by subtracting the averaged neighboring background fluorescence intensity lateral to the spindle, normalized to the area of the region of interest. Fluorescence signal of Cin8-3GFP, Bim1-GFP and Ase1-3GFP was quantified as above using SlideBook, RRID:SCR_014423. Kar3-3YFP fluorescence intensity along the spindle between the SPBs was calculated in ImageJ, RRID:SCR_003070, using a line-scan between the SPBs, excluding the signal peaks associated with the SPBs. Fluorescent intensity experiments and measurements were performed blinded. The plates were blinded before cells were cultured and all downstream treatments including scoring and intensity analysis were done on the blinded images. For figure display panels, corresponding fluorescence images were adjusted equally using SlideBook.

### Statistical analyses

Samples were based on cell genotype and did not require randomization. Unless otherwise noted, statistical significance was checked with an independent two-tailed t-test. Statistical analysis was performed using Microsoft Excel (RRID:SCR_016137) and GraphPad Prism, RRID:SCR_002798. To determine the start and end lengths of the initial fast phase of anaphase, linear regression analysis was performed using Prism, RRID:SCR_002798. Graphs were generated using GraphPad Prism. Error bars are defined in figure legends. SEM and SD refer to standard error of the mean and standard deviation, respectively. * p ≤ 0.05; ** p ≤ 0.01; *** p ≤ 0.001; n.s. = not statistically significant.

## Online Supplemental Material

This manuscript contains one supplementary figure and one supplementary table. Supplementary Figure S1 shows spindle localization of Cin8-3GFP, Kar3-3YFP, and Ase1-3GFP during the slow phase of anaphase, and Bim1-GFP during the fast and slow phases. Table S1 contains the yeast strains and plasmids used.

## Acknowledgments

The authors are grateful to W-L. Lee and J. Moore for useful reagents and thank F. Kiyimba for helpful feedback on the manuscript. This work was supported by an NSF grant (MCB-1846262) to MLG.

## Author Contributions

ETN and MLG designed the research; AB, ETN, LS, MLG, RMG, and VT constructed reagents, performed experiments and analyzed data; ETN and MLG wrote the manuscript; all authors participated in making figures and editing the manuscript.

## Contact for reagent and resource sharing

Additional information and requests for resources and reagents should be directed to and will be responded to by the Lead Contact, Mohan Gupta.

## Supplementary Figure Legends

**Supplementary Figure 1. Localization of MAPs to spindles composed exclusively of Tub1 or Tub3.**

(**a-d**) Ase1 localization to the spindle. (**a**) Representative DIC and wide-field fluorescence images of Ase1-3GFP localization to the spindle (microtubules) in Tub1-only (top) and Tub3-only (bottom) cells. Hashed box in merged represents area of spindle quantified. (**b-d**) Ase1-3GFP fluorescence intensity on spindles (**b**) in all anaphase cells, (**c**) during the initial, fast phase and (**d**) during the subsequent, slow phase of anaphase. (**e-g**) Bim1 localization to the spindle. (**e**) Representative DIC and wide-field fluorescence images of Bim1-GFP localization to the spindle (microtubules) in Tub1-only (top) and Tub3-only (bottom) cells. Hashed box in merged represents area of spindle quantified. (**f-g**) Bim1-GFP fluorescence intensity on spindles (**f**) during the initial, fast phase and (**g**) during the subsequent, slow phase of anaphase. (**h-i**) Fluorescence intensity of (**h**) Cin8-3GFP and (**i**) Kar3-3YFP on spindles during the slow phase of anaphase. Graphs show mean ± SEM. In (**b**) n for Tub1-only = 18, 25, 14, WT-Tub1 = 19, 23, 15, WT-Tub3 = 7, 13, 16, Tub3-only = 18, 10, 13, in (**c**) n for Tub1-only = 44, 46, 28, WT-Tub1 = 42, 43, 37, WT-Tub3 = 37, 41, 38, Tub3-only = 33, 33, 35, in (**d**) n for Tub1-only = 26, 21, 14, WT-Tub1 = 23, 20, 22, WT-Tub3 = 30, 28, 22, Tub3-only = 15, 23, 22 from three trials; in (**f**) n for Tub1-only = 17, 17, WT-Tub1 = 15, 13, WT-Tub3 = 15, 12, Tub3-only = 21, 15, and in (**g**) n for Tub1-only = 25, 23, WT-Tub1 = 24, 24, WT-Tub3 = 18, 21, Tub3-only = 23, 20 from two trials; and in (**h**) n for Tub1-only = 20, 21, 20, WT-Tub1 = 24, 23, 20, WT-Tub3 = 30, 34, 26, Tub3-only = 31, 26, 33, and in (**i**) n for Tub1-only = 11, 11, 10, WT-Tub1 = 10, 16, 10, WT-Tub3 = 10, 10, 19, Tub3-only = 12, 16, 10 from three trials. Scale bars = 2 μm.

