## Supplemental Figure 1 and Table 1 for "Tubulin isotypes contribute opposing properties to balance anaphase spindle morphogenesis"

### Figure S1

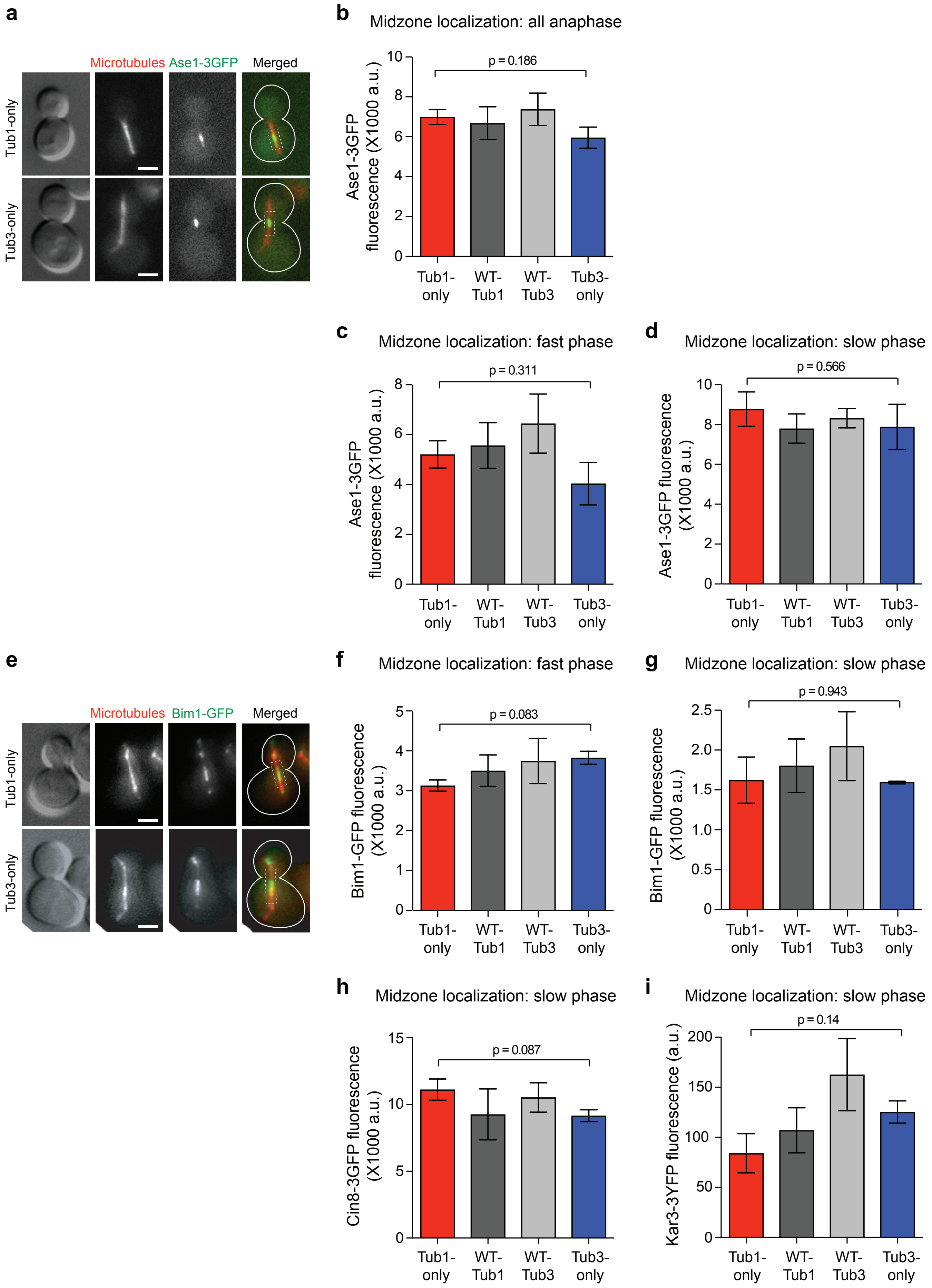

a
Microtubules
Ase1-3GFP
Merged
Tub1-only
Tub3-only
b
Midzone localization: all anaphase
Ase1-3GFP fluorescence (X1000 a.u.)
p = 0.186
0
2
4
6
8
10
Tub1-only
WT-Tub1
WT-Tub3
Tub3-only
c
Midzone localization: fast phase
Ase1-3GFP fluorescence (X1000 a.u.)
p = 0.311
0
2
4
6
8
Tub1-only
WT-Tub1
WT-Tub3
Tub3-only
d
Midzone localization: slow phase
Ase1-3GFP fluorescence (X1000 a.u.)
p = 0.566
0
2
4
6
8
10
Tub1-only
WT-Tub1
WT-Tub3
Tub3-only
e
Microtubules
Bim1-GFP
Merged
Tub1-only
Tub3-only
f
Midzone localization: fast phase
Bim1-GFP fluorescence (X1000 a.u.)
p = 0.083
0
1
2
3
4
5
Tub1-only
WT-Tub1
WT-Tub3
Tub3-only
g
Midzone localization: slow phase
Bim1-GFP fluorescence (X1000 a.u.)
p = 0.943
0
0.5
1.0
1.5
2.0
2.5
Tub1-only
WT-Tub1
WT-Tub3
Tub3-only
h
Midzone localization: slow phase
Cin8-3GFP fluorescence (X1000 a.u.)
p = 0.087
0
5
10
15
Tub1-only
WT-Tub1
WT-Tub3
Tub3-only
i
Midzone localization: slow phase
Kar3-3YFP fluorescence (a.u.)
p = 0.14
0
50
100
150
200
Tub1-only
WT-Tub1
WT-Tub3
Tub3-only

**Supplemental Table 1.** Yeast strains and plasmids used in this study.

| Strain name | Relevant genotype[a] | Source |
|---|---|---|
| MGY50 | MAT**a** ura3-52 his3-Δ200 leu2-Δ1 trp1-Δ63 | WT Control |
| MGY51 | MATα ura3-52 his3-Δ200 leu2-Δ1 trp1-Δ63 | WT Control |
| MGY2256 | MAT**a** ura3-52 his3-Δ200 leu2-Δ1 trp1-Δ63 tub3Δ::TUB1-NatMX4 | *Nsamba et al.* |
| MGY2257 | MAT**a** ura3-52 his3-Δ200 leu2-Δ1 trp1-Δ63 tub1Δ::TUB3-NatMX4 | *Nsamba et al.* |
| MGY2258 | MAT**a** his3-Δ200 leu2-Δ1 trp1-Δ63 pTUB1-GFP-TUB1-URA3 | *Nsamba et al.* |
| MGY2259 | MAT**a** his3-Δ200 leu2-Δ1 trp1-Δ63 pTUB1-GFP-TUB3-URA3 | *Nsamba et al.* |
| MGY2260 | MAT**a** his3-Δ200 leu2-Δ1 trp1-Δ63 tub3Δ::TUB1-NatMX4 pTUB1-GFP-TUB1-URA3 | *Nsamba et al.* |
| MGY2413 | MATα his3-Δ200 leu2-Δ1 trp1-Δ63 tub3Δ::TUB1-NatMX4 pTUB1-GFP-TUB1-URA3 | This study |
| MGY2261 | MAT**a** his3-Δ200 leu2-Δ1 trp1-Δ63 tub1Δ::TUB3-NatMX4 pTUB1-GFP-TUB3-URA3 | *Nsamba et al.* |
| MGY2414 | MAT**a** his3-Δ200 leu2-Δ1 trp1-Δ63 tub1Δ::TUB3-NatMX4 pTUB1-GFP-TUB3-URA3 | This study |
| MGY2317 | MAT**a** his3-Δ200 leu2-Δ1 trp1-Δ63 pTUB1-GFP-TUB1-URA3 CNM67-mRUBY2-G418 | This study |
| MGY2318 | MAT**a** his3-Δ200 leu2-Δ1 trp1-Δ63 pTUB1-GFP-TUB3-URA3 CNM67-mRUBY2-G418 | This study |
| MGY2319 | MAT**a** his3-Δ200 leu2-Δ1 trp1-Δ63 tub3Δ::TUB1-NatMX4 pTUB1-GFP-TUB1-URA3 CNM67-mRUBY2-G418 | This study |
| MGY2320 | MAT**a** his3-Δ200 leu2-Δ1 trp1-Δ63 tub1Δ::TUB3-NatMX4 pTUB1-GFP-TUB3-URA3 CNM67-mRUBY2-G418 | This study |
| MGY2321 | MATα his3-Δ200 leu2-Δ1 trp1-Δ63 pTUB1-mRUBY3-TUB1-URA3 KAR3-3YFP-LEU2 | This study |
| MGY2322 | MAT**a** his3-Δ200 leu2-Δ1 trp1-Δ63 pTUB1-mRUBY3-TUB3-URA3 KAR3-3YFP-LEU2 | This study |
| MGY2323 | MATα his3-Δ200 leu2-Δ1 trp1-Δ63 tub3Δ::TUB1-NatMX4 pTUB1-mRUBY3-TUB1-URA3 KAR3-3YFP-LEU2 | This study |
| MGY2324 | MATα his3-Δ200 leu2-Δ1 trp1-Δ63 tub1Δ::TUB3-NatMX4 pTUB1-mRUBY3-TUB3-URA3 KAR3-3YFP-LEU2 | This study |
| MGY2325 | MAT**a** his3-Δ200 leu2-Δ1 trp1-Δ63 pTUB1-mRUBY3-TUB1-URA3 CIN8-3GFP-TRP1 | This study |
| MGY2326 | MATα his3-Δ200 leu2-Δ1 trp1-Δ63 pTUB1-mRUBY3-TUB3-URA3 CIN8-3GFP-TRP1 | This study |
| MGY2327 | MATα his3-Δ200 leu2-Δ1 trp1-Δ63 tub3Δ::TUB1-NatMX4 pTUB1-mRUBY3-TUB1-URA3 CIN8-3GFP-TRP1 | This study |
| MGY2328 | MAT**a** his3-Δ200 leu2-Δ1 trp1-Δ63 tub1Δ::TUB3-NatMX4 pTUB1-mRUBY3-TUB3-URA3 CIN8-3GFP-TRP1 | This study |
| MGY2329 | MATα his3-Δ200 leu2-Δ1 trp1-Δ63 pTUB1-mRUBY3-TUB1-URA3 ASE1-3GFP-TRP1 | This study |
| MGY2330 | MAT**a** his3-Δ200 leu2-Δ1 trp1-Δ63 pTUB1-mRUBY3-TUB3-URA3 ASE1-3GFP-TRP1 | This study |
| MGY2331 | MATα his3-Δ200 leu2-Δ1 trp1-Δ63 tub3Δ::TUB1-NatMX4 pTUB1-mRUBY3-TUB1-URA3 ASE1-3GFP-TRP1 | This study |
| MGY2332 | MATα his3-Δ200 leu2-Δ1 trp1-Δ63 tub1Δ::TUB3-NatMX4 pTUB1-mRUBY3-TUB3-URA3 ASE1-3GFP-TRP1 | This study |
| MGY2282 | MAT**a** his3-Δ200 leu2-Δ1 trp1-Δ63 pTUB1-mRUBY3-TUB1-URA3 BIM1-GFP-HIS3 | *Nsamba et al.* |
| MGY2283 | MAT**a** his3-Δ200 leu2-Δ1 trp1-Δ63 pTUB1-mRUBY3-TUB3-URA3 BIM1-GFP-HIS3 | *Nsamba et al.* |
| MGY2284 | MAT**a** his3-Δ200 leu2-Δ1 trp1-Δ63 tub3Δ::TUB1-NatMX4 pTUB1-mRUBY3-TUB1-URA3 BIM1-GFP-HIS3 | *Nsamba et al.* |
| MGY2285 | MATα his3-Δ200 leu2-Δ1 trp1-Δ63 tub1Δ::TUB3-NatMX4 pTUB1-mRUBY3-TUB3-URA3 BIM1-GFP-HIS3 | *Nsamba et al.* |

| | | |
|---|---|---|
| MGY2415 | *MATα his3-Δ200 leu2-Δ1 dyn1Δ::TRP1 pTUB1-GFP-TUB1-URA3* | This study |
| MGY2245 | *MAT***a** *his3-Δ200 leu2-Δ1 dyn1Δ::TRP1 pTUB1-GFP-TUB1-URA3 BIK1-3GFP-NatMX4* | This study |
| MGY2369 | *MAT***a** *his3-Δ200 leu2-Δ1 dyn1Δ::TRP1 tub3Δ::TUB1-NatMX4 pTUB1-GFP-TUB1-URA3* | This study |
| MGY2416 | *MAT***a** *his3-Δ200 leu2-Δ1 dyn1Δ::TRP1 tub3Δ::TUB1-NatMX4 pTUB1-GFP-TUB1-URA3* | This study |
| MGY2398 | *MATα his3-Δ200 leu2-Δ1 dyn1Δ::TRP1 tub1Δ::TUB3-NatMX4 pTUB1-GFP-TUB3-URA3* | This study |
| MGY2399 | *MAT***a** *his3-Δ200 leu2-Δ1 dyn1Δ::TRP1 tub1Δ::TUB3-NatMX4 pTUB1-GFP-TUB3-URA3* | This study |
| MGY2372 | *MATα leu2-Δ1 trp1-Δ63 lys2-801 pTUB1-GFP-TUB1-URA3 she1Δ::HIS3* | This study |
| MGY2373 | *MAT***a** *leu2-Δ1 trp1-Δ63 lys2-801 tub3Δ::TUB1-NatMX4 pTUB1-GFP-TUB1-URA3 she1Δ::HIS3* | This study |
| MGY2374 | *MAT***a** *leu2-Δ1 lys2-801 pTUB1-GFP-TUB1-URA3 she1Δ::HIS3 dyn1Δ::TRP1* | This study |
| MGY2375 | *MATα leu2-Δ1 tub3Δ::TUB1-NatMX4 pTUB1-GFP-TUB1-URA3 she1Δ::HIS3 dyn1Δ::TRP1* | This study |

| **Plasmid name** | **Description and markers** | **Enzyme** | **Source** |
|---|---|---|---|
| pAFS125 | *pTUB1-GFP-TUB1-URA3*, *AmpR* (pMG210 in our records) | StuI | Straight *et al.* |
| pMG735 | *pTUB1-GFP-TUB3-URA3*, *AmpR*. Identical to pAFS125 but containing the *TUB3* ORF in place of *TUB1* ORF. | StuI | *Nsamba et al.* |
| pMG684 | *pTUB1-yomRUBY3-TUB1-URA3, AmpR* (yomRUBY3 = yeast-optimized monomeric RUBY3) | StuI | *Nsamba et al.* |
| pMG685 | *pTUB1-yomRUBY3-TUB3-URA3, AmpR.* Identical to pMG684 but containing the *TUB3* ORF in place of *TUB1* ORF. (yomRUBY3 = yeast-optimized monomeric RUBY3) | StuI | *Nsamba et al.* |
| pMG100 | *KAR3-C-terminus-3YFP-LEU2, AmpR.* Plasmid to tag endogenous KAR3 with *-3YFP*. | NheI | This study. |
| pJM0213 | *CIN8-C-terminus-3GFP-TRP1, AmpR.* C-terminal 670 bases of *CIN8* ORF was cloned into BamH1 of PB1586 (pRS304 with 3xGFP). Plasmid to tag endogenous *CIN8* with *-3GFP*. (pMG738 in our records). | EcoRI | Aiken *et al* |
| pMG701 | *ASE1-C-terminus-3GFP-TRP1, AmpR.* Plasmid to tag endogenous *ASE1* with *-3GFP*. | EcoRI | Zhu *et al.* |

[a]All yeast strains are of S288C background.

Aiken, J., D. Sept, M. Costanzo, C. Boone, J.A. Cooper, and J.K. Moore. 2014. Genome-wide analysis reveals novel and discrete functions for tubulin carboxy-terminal tails. *Curr. Biol.* 24:1295–1303. doi:10.1016/j.cub.2014.03.078.

Nsamba, E.T., A. Bera, M. Costanzo, C. Boone, and M.L. Gupta. 2021. Tubulin isotypes optimize distinct spindle positioning mechanisms during yeast mitosis. *J Cell Biol*. 220:e202010155. doi:10.1083/jcb.202010155.

Straight, A.F., Marshall, W.F., Sedat, JW & Murray, A.W. Mitosis in living budding yeast: anaphase A but no metaphase plate. *Science* **277**, 574–578 (1997).

Zhu, Y., X. An, A. Tomaszewski, P.K. Hepler, and W.-L. Lee. 2017. Microtubule crosslinking activity of She1 ensures spindle stability for spindle positioning. *J. Cell Biol.* 216:2759–2775. doi:10.1083/jcb.201701094
